# Functional recovery of the adult murine hippocampus after cryopreservation by vitrification

**DOI:** 10.1101/2025.01.22.634384

**Authors:** Alexander German, Enes Yağız Akdaş, Cassandra Flügel-Koch, Anna Fejtova, Jürgen Winkler, Christian Alzheimer, Fang Zheng

## Abstract

Cryopreserving the adult brain is challenging due to damage from ice formation, and traditional freezing methods fail to maintain neural architecture and function. Vitrification offers a promising alternative but has not been surveyed in the brain. Here, we demonstrate near-physiological recovery of the adult murine hippocampus after vitrification of brain slices and of the whole brain *in situ*. Key features of the hippocampus are preserved, including structural integrity, metabolic responsiveness, neuronal excitability, and synaptic transmission and plasticity. Notably, hippocampal long-term potentiation was well preserved, indicating that the cellular machinery of learning and memory remains operational. These findings extend known biophysical limits for cerebral hypothermic shutdown by demonstrating recovery after complete cessation of molecular mobility in the vitreous state. This suggests that the brain can be arrested in time and then reactivated, opening avenues for potential clinical applications.

**Significance Statement:** While the brain is considered exceptionally sensitive, we show that the hippocampus can resume normal electrophysiological activity after being rendered completely immobile in a cryogenic glass. The work extends known biophysical tolerance limits for the brain from the hypothermic to the cryogenic range and establishes a protocol for its long-term storage in a viable state.

## Introduction

Bridging space and time via life-suspending technologies is a familiar theme from fiction (1, 2), and it has found short-term application through procedures like deep hypothermic circulatory arrest (DHCA (3)). The brain’s remarkable capacity to recover from hypothermic shutdown demonstrates the potential to re-initialize brain dynamics (4), and supports the tenet that memory is structurally encoded in the synaptic architecture, a fundamental assumption in connectomics (5). However, it is not known whether brain function can be restored after complete shutdown of molecular mobility. Here, we establish near-physiological functional recovery after cryopreservation by vitrification of the hippocampus in adult mice. This has immediate applications: For example, it allows the distribution of neuroscientific experiments across different time points and locations, improving repro-ducibility and animal welfare (6). Furthermore, cryosubstitution of vitrified samples enables structural analysis in a near-native state, e.g., for connectomics (7, 8). Progress in cryopreservation of rodent organs has moved the theme of life-suspending technologies closer to plausibility (9-13), with the nervous system remaining as a cornerstone on the path towards clinical applications (14-16). Finally, empirical proof that brain function can be arrested in time could inform philosophical debates, e.g., whether mental phenomena resist physical explanation (17-19).

Already in 1953, Luyet and Gonzales demonstrated the possibility to cryopreserve embryonic avian brain tissue (20), and cellular survival after cryopreservation by freezing was later also shown for fetal rodent and human brain tissue (21-24). In the case of adult mammalian brain tissue and neurons differentiated in culture, full electrophysiological recovery after cryopreservation has thus far not succeeded (25-27). Using freezing techniques, some recovery of activity was reported in the adult feline brain after several years of storage at -20 °C (28, 29), and in rodent ganglia after up to 24 hours storage at -76 °C (30), both using 15% glycerol. Xue et al. showed recovery of neural organoids after freezing, using 10% dimethyl sulfoxide and 10% ethylene glycol combined with a ROCK-inhibitor (31). However, freezing of neural tissue results in overt loss of synaptic connections (32, 33). Achieving full electrophysiological recovery therefore likely necessitates ice-free cryopreservation, i.e., vitrification.

During cryopreservation by vitrification, the aqueous phase of biological tissue solidifies into a non-crystalline amorphous glass (34, 35). This can be achieved by replacing high proportions of the tissue water with polar solvents. Provided sufficient cooling and rewarming rates, these solvents act as cryoprotectants, inhibiting crystallization due to water-water interactions (36). Ice-free cryopreservation to avoid mechanical disruption of the inter- and intracellular space in complex biological materials has been proposed long ago (37, 38), and has been realized by vitrification (34, 39). Recent progress in rapid and uniform warming techniques has finally enabled cryopreservation by vitrification of the rat heart (10), liver (11) and kidney (9, 12, 13).

Pichugin et al. have demonstrated recovery of K^+^/Na^+^ ratios after vitrification of rat hippocampal slices at -130 °C (33). Building on this finding, we vitrified murine brain slices and the whole murine brain *in situ*, using a variant of the vitrification solution VM3 which we call V3 (40), containing a mixture of dimethyl sulfoxide, formamide and ethylene glycol. We held the specimens below the glass transition temperature for multiple days, ensuring complete cessation of molecular mobility. With an optimized vitrification procedure, after rewarming, we were able to achieve recovery of cellular metabolism and electrophysiology by patch-clamp recordings for individual cell activity and extracellular field potential recordings for short-term and long-term synaptic plasticity. This establishes a bona fide method to preserve functional brain tissue.

## Results

### Vitrification protocol for hippocampal slices

Based on stereomicroscopy assessment of tissue swelling and crystallization, as well as the degree of electrophysiological recovery, we optimized a vitrification procedure that minimizes damage from the following six variables (36, 40): 1. Toxicity of cryoprotective agents (CPA), determined by their composition, concentration, temperature and exposure time before vitrification and after rewarming (40). 2. Osmotic shrinking determined by the CPA loading protocol before vitrification (41-43). 3. Osmotic swelling determined by the CPA unloading protocol after rewarming (41-43). 4. Crystallization determined by the CPA concentration, composition, diffusion, and the rate of cooling for vitrification and the rate of rewarming (34). 5. Physical cracking determined by thermomechanical stress during cooling and rewarming in the vitreous state (44). 6. Chilling injury determined by temperature and exposure time before vitrification and after rewarming (45).

**Fig. 1A** illustrates our optimized vitrification procedure for adult murine brain slices. To minimize toxicity, osmotic shrinking, and chilling injury during CPA loading, slices were placed on a polyester mesh insert, and went through a protocol starting with CPA loading in LM5 carrier solution (containing 22.6 mM potassium, composition see **table S1**) containing 0%, 2%, 4%, 8%, 16% and 30% w/v vitrification solution V3 (**table S2**) at 10 °C. This was followed by 45% w/v V3 and three iterations of 59% w/v V3 at -10 °C. Reliability was increased by precise preparation of V3 using a volumetric flask and by consistently submerging the brain slices into the above solutions with a brush. Brain slices were directionally cooled from below, by placing them on a copper cylinder cooled to -196 °C with liquid nitrogen (**fig. S1**). The mesh insert carrying the slices was first transferred onto the top of the cylinder. After one minute, the mesh insert was slowly transferred into liquid nitrogen. This allowed us to avoid physical cracking of vitreous brain slices due to thermomechanical stress, as would occur during direct immersion into liquid nitrogen (**fig. S2**).

**Fig. 1.**
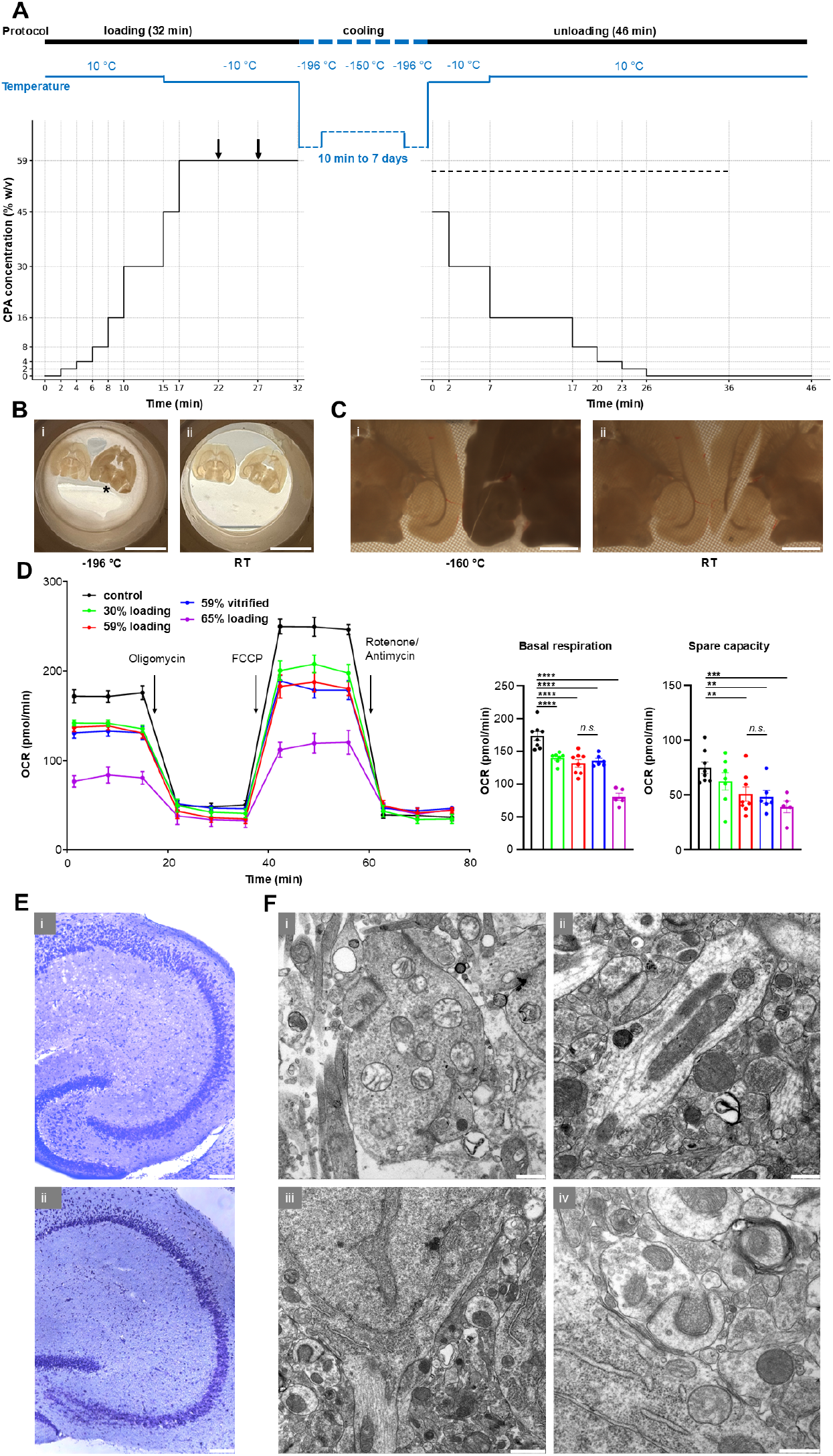
Vitrification of adult murine brain slices. (**A**) Time course of vitrification protocol, with varied temperature (**top**) and CPA concentration (**bottom**). Arrows indicate where transfer to fresh vitrification solution occurs. Dotted line indicates addition of 300 mM mannitol to the carrier solution for unloading. (**B**) Brain slices (350 µm thick) inside mesh at liquid nitrogen temperature (**i**; at -196 °C) demonstrating apparent vitrification (glossy and transparent) vs. crystallization (dull and opaque; asterisk). Such difference in appearance disappeared after the rewarming process (**ii**; 5 min exposure to RT). (**C**) Stereomicroscopy images of brain slices (350 µm thick) inside mesh covered with isopentane at isopentane melting point temperature (**i**; -160 °C) demonstrating apparent vitrification (transparent) vs. crystallization and cracking (opaque). Difference in appearance disappeared after the rewarming process, with the crack becoming clearly visible by separation of brain tissue (**ii**; 5 min exposure to RT). (**D**) Concentration-dependent effect of CPA loading on oxygen consumption rates (OCR) of murine hippocampal tissue. 0-20 min: Basal respiration. 20-40 min: Ablation of ATP-linked respiration with Oligomycin. 40-60 min: Maximal respiratory capacity induced with FCCP, 60-80 min: Non-mitochondrial respiration induced with rotenone and antimycin A. Note that the vitrification with 59% w/v V3 did not affect metabolism compared to incubation with 59% w/v V3 alone. Statistical comparisons were performed with one-way ANOVA followed by Fisher’s LSD, n.s. not significant, **p<0.005 ***p<0.0005 ****p<0.0001. (**E**) Nissl staining of hippocampal sections (30 µm) without treatment (control; **i**) and after vitrification (5 hours post-rewarming; **ii**) indicated the cytoarchitectural integrity. **(F**) Electron microscopy revealed well-preserved ultrastructure in the CA1 region of the vitrified hippocampus. Immediately after completion of the vitrification protocol and CPA unloading, occasional astroglial and mitochondrial swelling was observed (**i**). After 10 hours of incubation in aCSF at RT, the ultrastructure, including mitochondria and synapses (**ii**), dendrite and neuron (**iii**), synapse and myelin (**iv**) is indistinguishable from control slices. Scale bars: 1 cm (**B**); 1 mm (**C**) 100 µm (**E**); 0.5 µm (**i, ii, iv; F**), 1 µm (**iii; F**).

After 10 min to 7 days in liquid nitrogen and then in a -150 °C freezer, slices were rewarmed in 52% V3 at -10 °C. CPA unloading was performed in LM5 with additional 300 mM mannitol to reduce osmotic swelling. The solutions used were 45% and 30% w/v V3 at -10 °C, 16%, 8%, 4%, 2%, 0% w/v V3 at 10 °C. Unloading was concluded with a final step of LM5 without added mannitol (**Fig. 1A, table S3**).

The peak concentration of 59% w/v V3 was determined by varying the final CPA loading step (at -10 °C) with concentrations ranging from 45% to 65% w/v V3, as vitrification properties are tissue dependent and empirical (46). During cooling and rewarming, successful cryopreservation was predictable by direct visual observation, with vitreous slices remaining glossy and transparent (**Fig. 1B and C**). Failure of cryopreservation was predictable when slices turned dull and opaque, due to apparent cooling phase crystallization (**Fig. 1B and C**) and/or transient rewarming phase crystallization ((47), **fig. S3, Movie S1-S7**). Using the stereomicroscope, we observed complete avoidance of cooling phase crystallization with CPA concentrations starting from 58%. Complete avoidance of transient rewarming phase crystallization was observed starting from 56% w/v V3 when using rapid rewarming (**Movie S8-S11**). Successful recovery after cryopreservation was achieved starting from 57% w/v V3. We observed complete avoidance of rewarming phase crystallization with 65% w/v V3 even during slow rewarming (**fig. S4, Movie S12**). The determining factor for brain slice vitrification in the final CPA loading step was therefore cooling phase crystallization. If not otherwise specified, for all presented analyses, a standard concentration of 59% w/v V3 was used to ensure complete avoidance of ice formation when using rapid rewarming.

Our optimized protocol had to balance damage from osmotic changes of cell volume due to transmembrane solute-solvent flux during concentration changes of CPA with the temperature- and time-dependent toxicity of higher CPA concentrations (41, 42). Limited cell shrinking and swelling can be beneficial to minimize intracellular exposure to peak CPA concentrations (39, 43). We varied concentration and timing of CPA loading and unloading steps. For example, all slices incubated in concentrations of 30% w/v V3 and above showed a transient swelling (8% increase in area) with faint whitening and loss of transparency during CPA unloading (**fig. S5**). This swelling appeared at 16% w/v V3 concentration at 10 °C and became more pronounced during incubation in decreasing concentration steps. It disappeared after 15 minutes of post-incubation in artificial cerebrospinal fluid (aCSF) at room temperature (RT). This transient swelling could be mitigated by extending the incubation time in 16% w/v V3 during unloading, improving subsequent electrophysiological recovery. Introducing an intermediate step of 45% w/v V3 during loading and unloading increased the yield of superficial neurons in subsequent patch-clamp recordings.

### Metabolic recovery

Mitochondrial activity of brain slices was evaluated in the CA1 region based on oxygen consumption rate (OCR, measured in pmol/min) using a previously established protocol (48). Using the optimized protocol, slices were loaded up to different peak CPA concentrations, subsequently unloaded and incubated for two hours in aCSF at room temperature (RT) for recovery. Cooling below -10 °C was omitted to enable comparison with vitrification. Peak CPA concentrations were: low (30% w/v V3 equivalent to 4.28 M permeable CPA, n = 7), standard (59% w/v V3 equivalent to 8.42 M permeable CPA, n = 6) and high (65% w/v V3 equivalent to 9.28 M permeable CPA, n = 5). These were compared to fresh tissue (control, n = 8) and vitrification with standard CPA concentration (n = 8) (**Fig. 1D**). CPA exposure reduced the basal respiration (control: 173.3 ± 6.7, low CPA: 139.7 ± 3.1, standard CPA: 135.1 ± 6.7, high CPA: 80.4 ± 5.6, Spearman’s r = −0.9018, permutation-based p = 0.0002). We did not observe a change between standard CPA and vitrification (standard CPA: 135.1 ± 6.7, vitrification: 131.4 ± 5.7). Higher concentrations of CPA significantly reduced the spare capacity as calculated in (48) (control: 73.4 ± 5.7, low CPA: 62.3 ± 8.0, standard CPA: 48.6 ± 5.8, high CPA: 38.9 ± 5.6, Spearman’s r = −0.6731, permutation-based p = 0.0004). Again, we did not observe a change between standard CPA and vitrification (standard CPA: 48.6 ± 5.8, vitrification: 50.6 ± 6.4). The metabolic analysis indicated a modest reduction in basal respiration after 30% w/v and 59% CPA loading, and markedly reduced rates after exposure to 65% w/v V3 solution to about half of control values. This sharply increasing toxicity of CPA was also manifested in the maximal respiratory capacity (**Fig. 1D**).

### Morphological assessment

Light and electron microscopic analyses were further employed to evaluate the potential morphological changes after standard CPA loading and vitrification. Nissl staining of brain sections (30 µm) revealed overall normal morphology after vitrification (n = 6 for 1-5 hours), compared to control untreated slices (**Fig. 1E)**. Electron microscopy of ultrathin sections (50 nm; **Fig. 1F**) indicated clear membranes of intact neuronal and synaptic structures in the hippocampus. Fixation performed immediately after completion of CPA unloading revealed limited mitochondrial swelling (**Fig. 1F(i)**). Like macroscopic swelling (**fig. S5**), ultrastructural swelling of this organelle resolved during incubation in aCSF at RT and was not observed in slices incubated for 1-10 hours after rewarming, which exhibited regular ultrastructure like untreated controls (n=4 for 1-10 hours, **Fig. 1F(ii)-(iv)**).

### Electrophysiological recovery after vitrification of brain slices

Next, we interrogated the functionality of the hippocampal circuitry after cryopreservation by vitrification with standard CPA concentration (59% w/v V3), focusing first on the Schaffer collateral (SC) - CA1 pyramidal cell synapse, arguably the best studied synapse in the mammalian brain. Combined with extracellular recording in CA1 stratum radiatum, electrical stimulation of SC evoked field postsynaptic potentials (fPSPs, **Fig. 2)** predominantly mediated by AMPA-type glutamate receptors, as indicated by their sensitivity to CNQX (40 µM; **fig. S6A**). When comparing untreated slices (n = 9 from 7 mice) with vitrified slices (n = 23 from 12 mice), the input-output (I-O) curve from the latter preparation showed a trend towards a lower gain in fPSP amplitude with increasing stimulus intensities (50 - 200 µA), leading to a downward deflection of the curve, which reached significance at the maximum response (**Fig. 2A**). Loading with 59% w/v CPA alone (n = 7 from 4 mice) produced an I-O curve that was situated in between the curves from untreated and vitrified slices (**Fig. 2A**, blue curve), without being significantly different from untreated controls. This finding suggests that, despite the known dose-dependent toxicity of CPA (40), the standard concentration used here for vitrification did not inflict appreciable damage upon basic synaptic properties. Note that even after loading with a substantially lower CPA concentration (30% w/v), the I-O curve tended to level off at high stimulus intensity (**fig. S6B**).

**Fig. 2.**
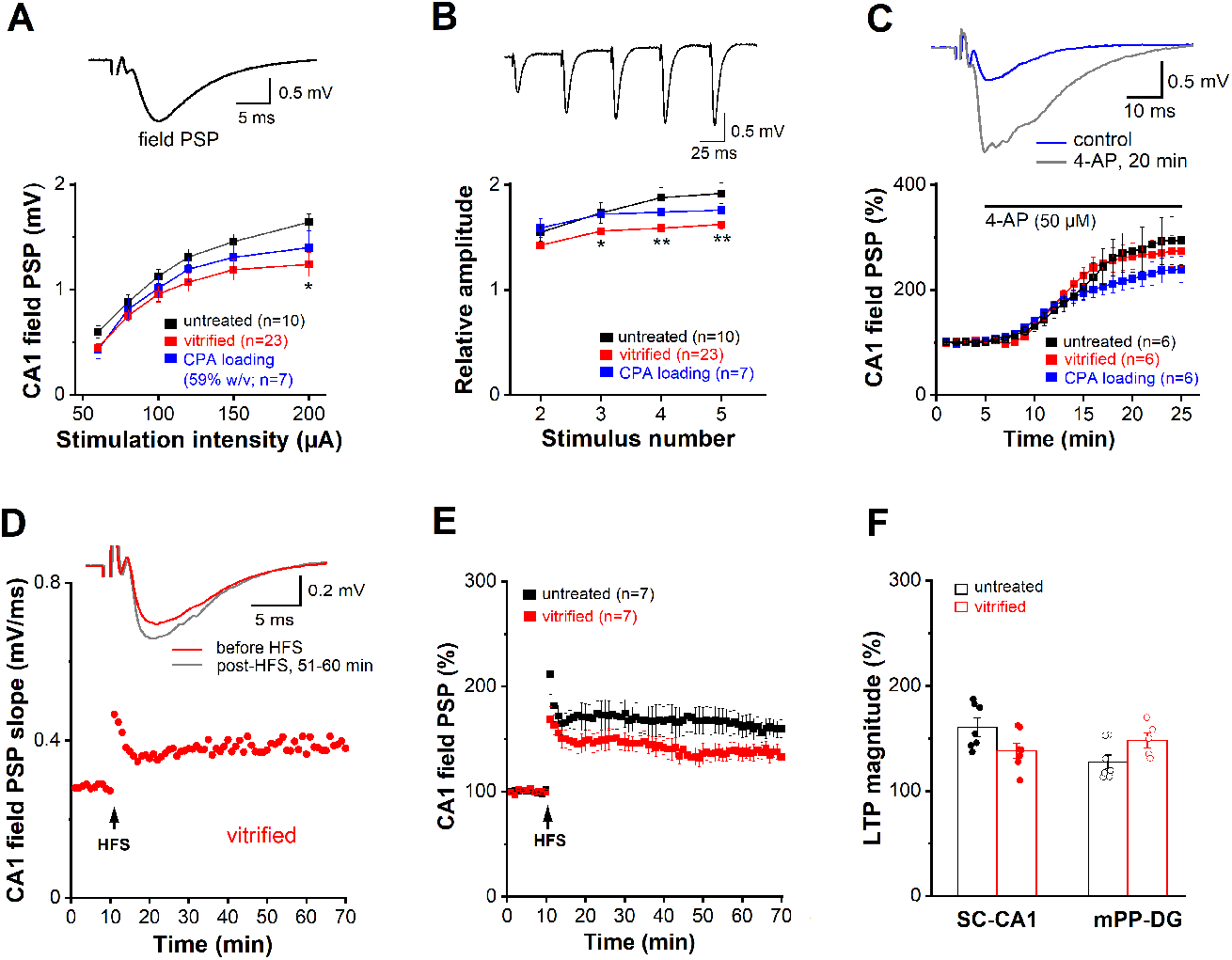
Recovery of synaptic functions after slice vitrification and rewarming. Field postsynaptic potentials (field PSP or fPSP) were evoked by electrical stimulation of Schaffer collaterals (SC) and monitored in CA1 stratum radiatum. **(A)** Input-output (I-O) curves from untreated slices, CPA loading controls, and vitrified slices. The inset on top illustrates typical field PSP response in an untreated slice (stimulus strength 75 µA). (**B**) Effects of vitrification and CPA loading alone on shortterm synaptic plasticity (STP). Same groups as in (**A**). Inset on top illustrates characteristic facilitation of fPSPs during STP-inducing stimulation (untreated slice, 70 µA). (**C**) Time course of fPSP augmentation during bath application of 4-AP (50 µM) in untreated slices, loading controls, and vitrified slices. Inset on top depicts effect of 4-AP on fPSP in slice with CPA loading alone. (**D-E**) LTP of SC-CA1 synapses in untreated controls and vitrified slices. Inset in (**D**) illustrates averaged voltage traces before and 51-60 min post high frequency stimulation (HFS). **(F)** Histogram depicts LTP magnitude as percentage increase in field PSP slope relative to baseline before HFS at the SC-CA1 and the mPP-GC synapse. Statistical comparisons were performed with one-way ANOVA followed by Tukey’s post-hoc test or Student t-test. * p < 0.05; ** p < 0.01.

Like many other excitatory synapses in the brain, the SC-CA1 synapse exhibits a rich repertoire of brief changes and persistent alterations, critically depending on the particular kind of input pattern. Whereas short-term plasticity (STP) involves mainly presynaptic mechanisms, long-term plasticity relies heavily on the postsynaptic machinery. We probed STP using a brief stimulus train (5 pulses at 20 Hz), with the stimulus intensity set to about 35% of that evoking the maximum response. We observed frequency facilitation in all groups (**Fig. 2B**). For the analysis, fPSP responses to 2^nd^ – 5^th^ stimuli were normalized to 1^st^ stimulus in the train. STP was significantly attenuated in vitrified slices, whereas loading alone did not alter the extent of STP measured in untreated slices (**Fig. 2B**). To exclude that an overall decrease in the transmitter pools that can be released and mobilized accounts for reduced STP after vitrification, we used the K^+^ channel blocker 4-aminopyridine (4-AP, 50 µM) to achieve maximum transmitter release from the terminals. When we plotted the gradual increase in fPSP amplitude as 4-AP was washed into the slice, neither the time course of drug action nor the maximum response that was eventually attained differed between the three groups (**Fig. 2C**).

We then examined long-term potentiation (LTP), the most important form of lasting plasticity and widely recognized as a key neurobiological substrate of learning and memory formation. Attesting to the remarkable tolerability of vitrification, cryopreserved hippocampal slices reliably produced LTP, which we induced using a standard protocol of high frequency stimulation (HFS, 100 Hz for 1 s, delivered twice 20 s apart **Figs. 2D-F**). Although the trajectories of potentiation tended to be different, when quantified over 51-60 min post-HFS, SC-CA1 LTP amounted to 160.50 ± 8.95 % in untreated slices (n = 7 from 7 mice), which was not significantly different from the extent of LTP in vitrified slices (138.06 ± 6.90 %, n = 7 from 7 mice; p = 0.070). To add weight to the striking finding of operational LTP post-vitrification, we extended our study to another essential synapse within the hippocampal formation that gates and processes incoming signals from the neocortex. This synapse is formed by the medial perforant path (mPP) of entorhinal cortex origin projecting onto granule cells (GC) of the dentate gyrus. Compared to untreated controls, the mPP-GC synapse in vitrified slices showed a downward shift of the I-O curve, especially across lower stimulation intensities (**fig. S6C**, left; untreated, n = 14 from 11 mice; vitrified, n = 9 from 4 mice). However, STP and LTP of mPP-GC synapses were not reduced in vitrified slices, indicating recovery of both transient and persistent synaptic plasticity after vitrification (**fig. S6C**, right**; Fig. 2F**)

To examine the effects of cryopreservation on cellular excitability, we performed whole-cell recordings from CA1 pyramidal cells (PCs) and dentate gyrus GCs in control and vitrified brain slices (with 59% w/v CPA) containing the dorsal hippocampus. Some cryopreserved cells were filled with biocytin to demonstrate the characteristic morphology with soma, apical and basal dendrites (**Fig. 3A**). In current-clamp mode, vitrified CA1 PCs had a resting membrane potential (RMP) of -74.82 ± 1.12 mV (n = 13 from 6 mice), which was virtually identical to that of untreated cells (−74.65 ± 0.73 mV; n = 21 from 10 mice, p = 0.779). The same was true for membrane capacitance (C_m_) and input resistance (R_m_) (**fig. S7A-C**). In contrast, vitrified GCs in dentate gyrus exhibited a small but significant depolarization of their RMP (vitrified-GC -83.53 ± 1.24 mV, n = 16 from 6 mice; control-GC -86.00 ± 0.45 mV, n = 29 from 17 mice; p = 0.023), accompanied by reduced C_m_ (vitrified GC 58.18 ± 3.17 pF; control GC 77.95 ± 2.88 pF; p = 0.0002) (**fig. S7A-B**). Lending further support to our finding of retained synaptic functionality in field potential recordings from cryopreserved slices (**Fig. 2**), spontaneous synaptic events were recorded regularly from both vitrified PCs and GCs. Moreover, the events occurred at the same frequency that was measured in their control counterparts (**Fig. 3B)**.

**Fig. 3.**
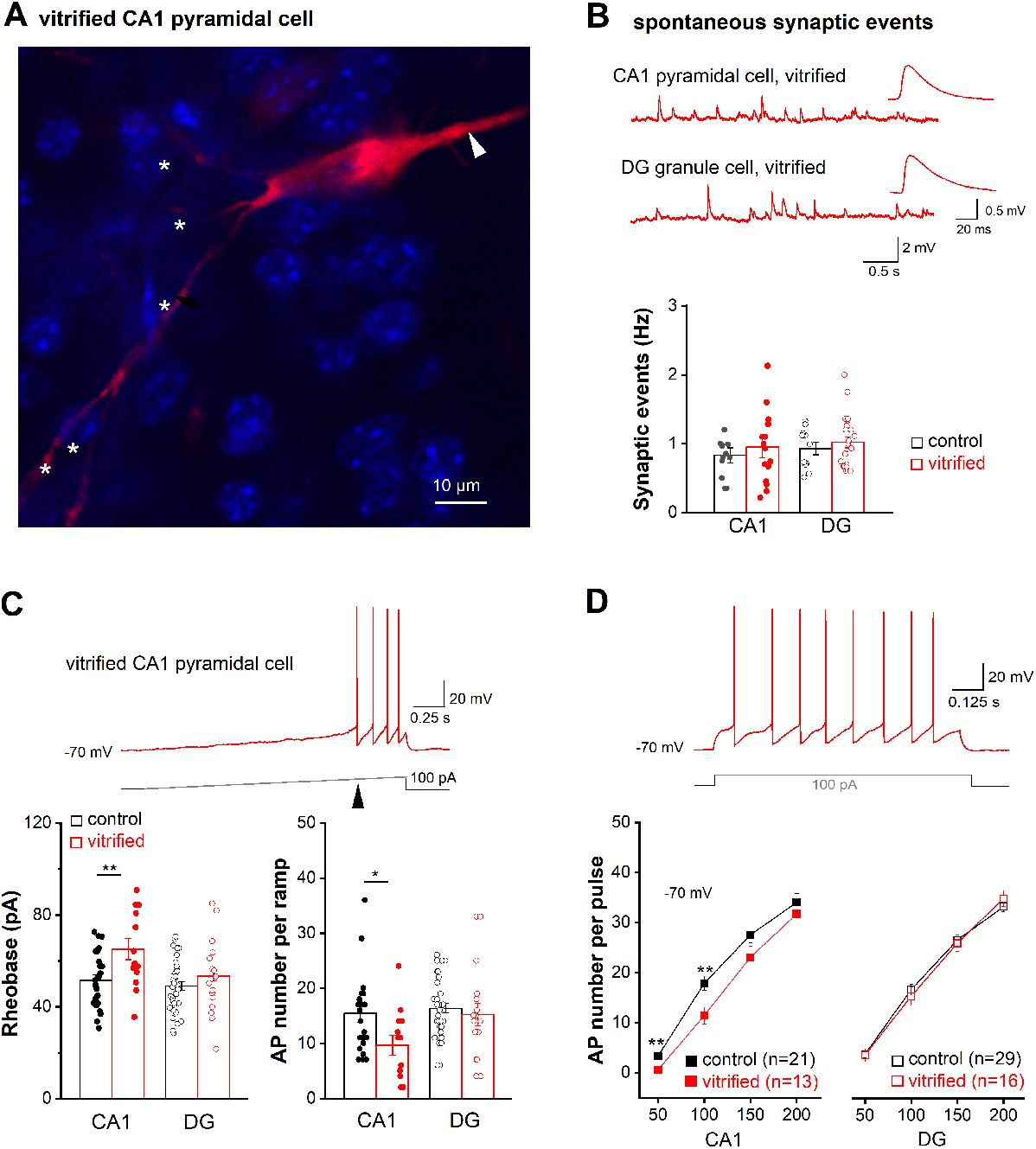
Recovery of cellular excitability and spontaneous synaptic activity after slice vitrification and rewarming. Whole-cell current-clamp recordings were performed from CA1 pyramidal cells and dentate gyrus (DG) granule cells.**(A)** Biocytin-filled CA1 pyramidal cell from a vitrified slice was stained with Cy3-coupled streptavidin (red). Arrowhead and asterisks point to apical dendrite and basal dendrites, respectively. Nuclei were stained with DAPI (blue). **(B)** Frequency of spontaneous synaptic events in CA1 and DG cells did not differ between control and vitrified slices. The traces above the histogram exemplify events in individual recordings taken at resting membrane potential. Insets depict averaged waveforms of spontaneous synaptic events at higher magnification. (**C-D**) Action potential (AP) firing was elicited using depolarizing currents delivered as ramps (C) or rectangular steps (D) from membrane potential adjusted to -70 mV. Arrowhead in current trace of (C) indicates rheobase. Histograms show that vitrification altered firing properties only in CA1 pyramidal cells, but not in DG granule cells. Statistical comparisons were performed using unpaired, two-tailed student’s t-test (**B, C**) or one-way ANOVA followed by Tukey’s post-hoc test (**D**). * p < 0.05; ** p < 0.01.

To probe firing properties, cells were held at -70 mV by DC injection before ramp-like or rectangular depolarizing current pulses were delivered. Ramp depolarization (0 to 100 pA within 2 s) revealed a marked increase in the current threshold evoking the first AP (i.e. rheobase) in vitrified PCs, whereas no such change was seen in vitrified GCs (**Fig. 3C**). Elevated rheobase in CA1 PCs was accompanied by reduced AP firing during ramps (**Fig. 3C**). When tested with rectangular depolarizing steps 1 s long, decreased firing in vitrified PCs became evident at smaller depolarizations (50 and 100 pA), but disappeared with larger steps (150 and 200 pA, **Fig. 3D**). Closer examination of AP waveforms from vitrified PCs revealed enhanced discharge threshold and increased afterhyperpolarization (AHP) as two potential mechanisms rendering these neurons less excitable than their untreated counterparts (**fig. S7E-I**). As vitrification did not impair hippocampal GC discharge pattern, we were surprised to find their AP waveforms differed from untreated controls in several respects including decreased threshold, faster rise time and smaller half-width (**fig. S7E-I**). Together with the reduced membrane capacitance, which makes the cell electrically more compact, these changes might result from reduced metabolic capacity, or reflect an adaptive process to reinstitute regular GC firing after vitrification (**Fig. 3C-D**).

The absence of neurophysiological signs of hyperexcitability, let alone seizure-like phenomena, in our field potential and whole-cell recordings from two distinct hippocampal regions strongly argues for an intact inhibitory network after vitrification. To substantiate this notion, we made whole-cell recordings from interneurons in the CA1 pyramidal cell layer of cryopreserved slices (n = 12 interneurons from 7 mice; **fig. S8**). These interneurons were characterized by low capacitance (26.67 ± 1.92 pF) and high input resistance (767.08 ± 156.86 MΩ). At rest, they were either quiet with an RMP of -67.56 ± 1.97 mV (n = 5; **fig. S8A**), or fired action potentials tonically (n = 5) or in bursts (n = 2) (**fig. S8C-D**). When held at -70 mV, the silent interneurons discharged 31.11 ± 8.86 APs (n = 5) in response to a 50 pA depolarizing current injection (for 1 s) (**fig. S8B**).

### Cerebral vitrification *in situ*

Successful cryopreservation of brain tissue slices encouraged us to scale this approach to the whole murine brain. However, initial trials with gradual CPA concentration steps (as employed for slices in **Fig. 1A**) during transaortic vascular perfusion led to extensive cerebral dehydration and macroscopic darkening (**Fig. 4Ai**), with reduction in cerebral mass to 44.5 ± 1.8% (N = 2) of control brains perfused with phosphate-buffered saline (PBS). Brains exposed to gradual CPA loading were macroscopically dehydrated to the extent that they became concave, with no fPSP response elicitable. Such dehydration might be attributed to a selectively high hydraulic conductivity within the blood-brain barrier (BBB). Shock-loading via perfusion with full-strength vitrification solution (59% w/v V3) improved total cerebral mass retention to 55.7 ± 3.0% (N = 5) of control. To harness the mismatch between the permeability for CPA and water within the BBB, we performed interleaved equilibration by alternating perfusion between full-strength vitrification solution and LM5 to rehydrate the partly CPA-loaded brain. Repeated cycles of interleaved equilibration enabled vitrification at normal cerebral mass and volume **(Fig. 4Aiii)**, but resulted in cerebral edema during CPA washout (**Fig. 4Aiv**). We therefore chose a final interleaved equilibration protocol that resulted in intermediate cerebral dehydration that maintained brain convexity (**Fig. 4Aii**), corresponding to a cerebral mass of 69.8 ± 13.5% (N = 5) of controls. Detailed information regarding the protocol for whole-brain cryopreservation is provided in the supplement and involves a craniectomy (**table S4; fig. S10, S11**). The brains remained *in situ* and were stored at -140°C for 1-8 days. After rewarming, CPA was washed out via transaortic vascular perfusion at reduced flow rates using a hyperoncotic solution (**table S4; fig. S9, S10, S11**).

**Fig. 4.**
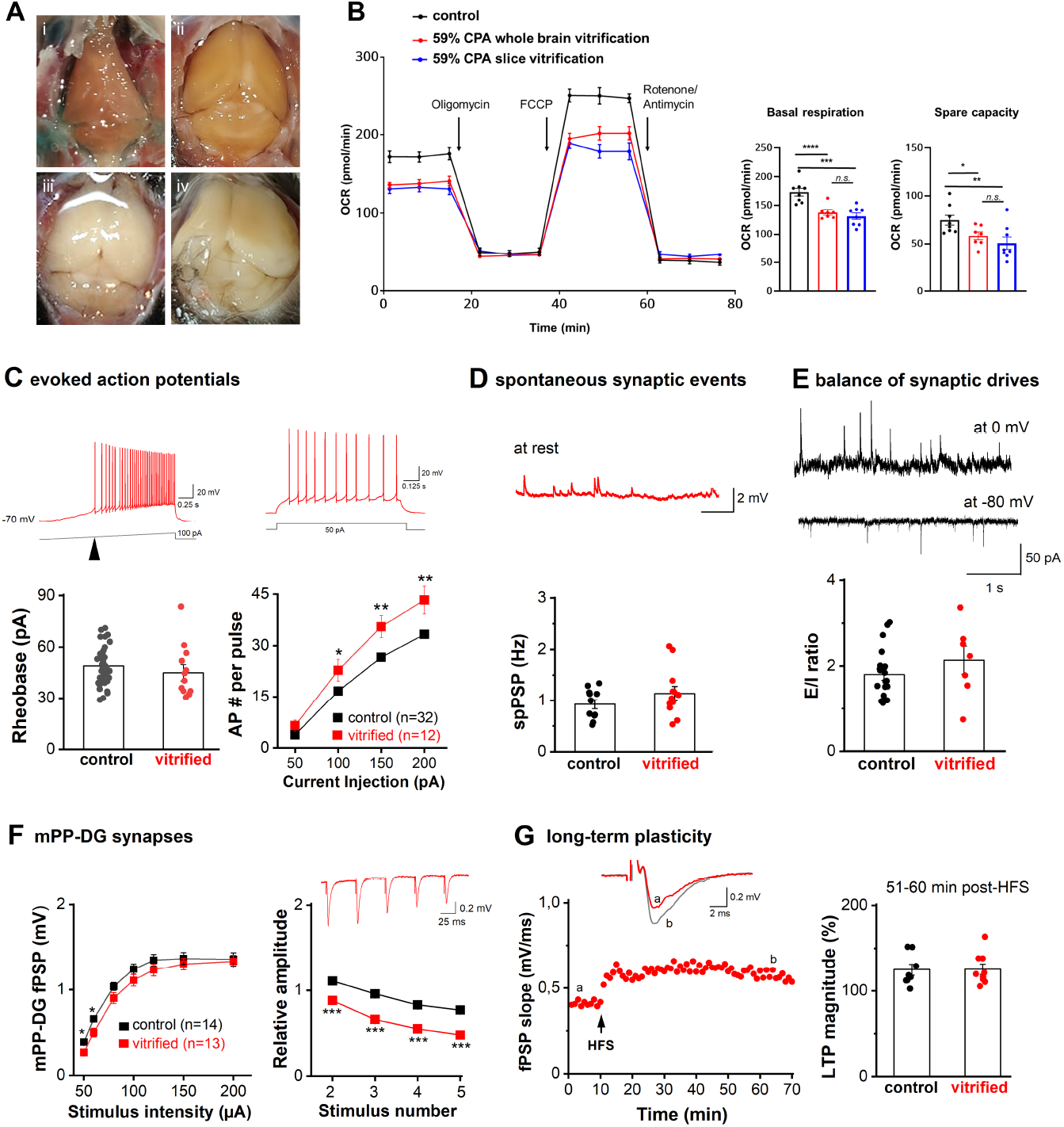
Hippocampal cellular excitability and synaptic function after whole brain CPA perfusion, vitrification and hyperoncotic washout *in situ*. **(A)** Photographs of the murine brain *in situ*, showing lethal cerebral dehydration after gradual CPA equilibration (**i**), intermediate volume reduction after successful interleaved equilibration (**ii**), insufficient volume reduction after interleaved equilibration (**iii**), and lethal cerebral edema after gradual CPA washout instead of hyperoncotic washout (**iv**). Functional evaluations in (**B-G**) were performed in acute hippocampal slices prepared from adult murine brains after vitrification in condition (**Aii**), storage at -140 °C for 1-8 days, and rewarming followed by hyperoncotic washout. **(B)** Mitochondrial activity assessed using a Seahorse analyzer across defined respiratory states: basal (0–20 min), oligomycin-inhibited (20–40 min), FCCP-stimulated (40–60 min), and rotenone/antimycin A-induced non-mitochondrial respiration (60–80 minHippocampi recovered from whole-brain vitrification with 59% w/v CPA showed a significant reduction in OCRs compared to controls. Data from the vitrified hippocampal slices were replicated here for comparison. Statistical analysis: one-way ANOVA with Fisher’s LSD; n.s. not significant, *p < 0.05, **p < 0.005, ***p < 0.0005, ****p < 0.0001. (**C**) Whole-cell current-clamp recordings of hippocampal granule cells in slices from the vitrified brains show comparable rheobase (indicated with arrowhead in current trace above) and enhanced excitability. As illustrated from voltage traces from a vitrified cell, depolarizing currents were delivered as ramp (left) or rectangular steps (right) from membrane potential adjusted to -70 mV. (**D**) Occurrence of spontaneous postsynaptic potentials (spPSPs), monitored at resting membrane potential under whole-cell current-clamp mode, did not differ between control and vitrified brains. (**E**) Whole-cell voltage-clamp recordings were performed to assess the balance of excitatory and inhibitory drives (E/I balance) onto untreated and vitrified DG cells. Membrane potential of DG granule cells was clamped either at -80 mV to monitor excitatory postsynaptic currents (EPSCs) or at 0 mV for inhibitory PSC (IPSC), as shown on the left with traces from a control slice. Histogram with E/I ratio, calculated with the frequency of PSCs in individual cells, indicated no appreciable alteration after cryopreservation. **(F-G)** mPP-DG granule cell synaptic transmission and plasticity after vitrification with interleaved equilibration. Compared to untreated controls, extracellularly recorded responses of mPP-DG synapses to electrical stimulation show well-preserved I-O curve for basic synaptic transmission after vitrification and rewarming (**F**, left panel). In contrast to significant reduction in STP (upon 20 Hz train stimuli; **F**, right panel), HFS-induced LTP of mPP-DG granule cell synapses was not affected after vitrification, as shown by example of time course for field PSP from a vitrified brain (**G**, left) and group data of normalized magnitude of change 51-60 min post tetanus (**G**, right). Statistical comparisons were performed using unpaired, two-tailed student’s t-test or one-way ANOVA followed by Tukey’s post-hoc test. * p < 0.05; ** p < 0.01.

One out of three iterations of the final protocol of interleaved equilibration, vitrification, and hyperoncotic washout yielded brain slices suitable for physiological evaluation in the Seahorse assay (N = 3, **Fig. 4**) and electrophysiological recordings (N = 10; **Fig. 4**). Tissue from these iterations demonstrated viability and physiological function after cryopreservation. The remaining iterations had to be excluded from the analysis due to either inadequate perfusion quality or absence of detectable electrophysiological activity.

We compared basal respiration and spare respiratory capacity of the hippocampi after whole brain vitrification (138.2 ± 4.45 pmol/min and 58.4 ± 3.99 pmol/min, respectively) to those of slice vitrification, and found no significant differences between the two conditions (**Fig. 4B**).

We then focused our electrophysiological experiments on hippocampal granule cells, given their observed viability in cryopreserved slices. We performed whole-cell recordings in current-clamp mode and in voltage-clamp mode to characterize their intrinsic excitability and the excitatory and inhibitory synaptic drives, respectively. Compared to untreated cells, DG granule cells in acute slices prepared from vitrified brains had comparable RMP (−85.17 ± 1.52 mV, n = 12 from 6 mice), input resistance (at -70 mV, 294.62 ± 18.44 MΩ), and rheobase (45.95 ± 4.46 pA) with evoked action potentials via ramp-like depolarization (**Fig. 4C**, left), however smaller Cm (62.23 ± 0.52 mpF; p = 0.025 vs control) as that observed from vitrified slices (**s. fig. 7B**). When rectangular depolarization (1 s) was delivered at larger steps (100 - 200 pA), increased discharge in GCs from vitrified brains was observed (**Fig. 4C**, right), suggesting an elevation in intrinsic excitability.

As observed for the slice protocol (**Fig. 3**), the functional synaptic network was manifested in spontaneously occurring postsynaptic potential (spPSP, 1.14 ± 0.14 Hz, n =12 from 6 mice; **Fig. 4D**). When the recordings were switched to voltage-clamp mode, glutamatergic excitatory drive and GABAergic inhibitory drive could be distinguished with voltage clamped at -80 mV and 0 mV, respectively (**Fig. 4E**). Such simultaneous monitoring enables a measurement of the overall balance between excitation and inhibition (E/I) in each cell, expressed as E/I ratio using the frequency of postsynaptic currents (PSCs). The E/I balance was unchanged in vitrified brains (vitrified 2.14 ± 0.33, n = 7 from 3 mice; untreated 1.80 ± 0.0.13, n = 19 from 4 mice; p = 0.255; **Fig. 4E**). In field potential recordings from mPP-GC synapses (n = 13 from 6 mice; **Fig. 4F-G**), basic synaptic transmission to mPP activation (I-O curve; **Fig. 4F, left**) was affected only at low stimulus intensity. STP was markedly reduced in DG of vitrified brains (**Fig. 4F, right**), which was in contrast to intact STP in vitrified slices (**s.fig.6C**), LTP of the mPP-GC synapse in vitrified brains was operational (51-60 min post-HFS, 125.39 ± 6.10 %, n = 9 from 4 mice; p = 0.913 vs untreated; **Fig. 4G**).

## Discussion

Thermal conditions must be carefully controlled for electrophysiological recordings, and brain tissue can become permanently unresponsive when temperatures exceed 42 °C (49). It has long been established that despite these sensitivities, the brain tolerates intermittent hypothermic-ischemic conditions remarkably well (50). Our results indicate that these tolerance limits extend to the cryogenic range, by demonstrating the functional recovery of the adult murine hippocampus after cryopreservation by vitrification and rewarming with retained morphological integrity, mitochondrial respiration, and electrophysiological activity, including synaptic transmission and plasticity. Normal spontaneous synaptic events revealed that brain activity re-initializes after cessation of all continuous dynamical processes in the vitreous state. Our work improves substantially upon previous attempts at cryopreserving adult brain tissue (28, 29, 33), in that vitrification was well tolerated in the short term, enabling near-physiological recovery of the various indices of intact synaptic transmission and cellular excitability. Previous electrophysiological recordings in neural tissue have focused on freezing (28, 29, 31), accompanied by a loss of synaptic connections (32). Using vitrification, not only was neural excitability recovered, but also synaptic transmission with the induction of robust LTP. Of note, the slices did tolerate a minimal degree of cooling phase crystallization (**Movie S9**). While vitrification has been applied successfully to peripheral organs (11-13), our study encourages its application in the central nervous system.

Notwithstanding the exceptional reinstitution of normal hippocampal functionality in the cryopreserved slices, we noted some differences in principal neurons from the CA1 region and the dentate gyrus weathered vitrification. While pyramidal cells from CA1 exhibited reduced excitability as indicated by increased rheobase and reduced AP firing, GCs from the dentate gyrus maintained normal discharge properties post-vitrification. The different sensitivity to the vitrification process might be due to variations in cell size, membrane properties, or intrinsic metabolic activity, as such variations could in turn result in different osmotic behaviour during CPA loading and unloading as well as in different susceptibility to direct CPA toxicity and chilling injury, e.g., by cold denaturation (45). The unchanged E/I balance in vitrified brains suggests that vitrification is unlikely to produce a pathological distortion of synaptic weights.

The peak concentration of V3 induced a dose-dependent reduction of the basal respiration and maximal respiratory capacity, indicating peak cryoprotectant toxicity as a cause of mitochondrial stress. Transient mitochondrial stress after vitrification was also implied by ultrastructural swelling of this organelle in electron microscopy. Of note, comparative biology suggests mitochondria play an important role in the adaptation to cold, e.g., in the canadian wood frog (51), or in insects (52). NMDA receptors have been shown to be sensitive to chilling (53), yet both SC-CA1 synapses and mPP-DG synapses, which express NMDA receptor-dependent LTP, showed reliable synaptic potentiation post-vitrification. Understanding these differences between cell types and between vitrified and untreated controls seems difficult, as hypothermia alone affects nearly every investigated biological pathway (50). It is possible that hypothermia during slicing combined with a brief rewarming and 30 min recovery phase at RT confers some protective preconditioning due to a cold-shock response (50).

The brain is sensitively dependent on homeostasis of its solute-solvent milieu to allow efficient neural computation. For example, potassium and water content of the cerebral extracellular space are tightly regulated and isolated from the blood compartment (54). Delivery of CPAs to the mammalian brain via perfusion is complicated by a mismatch between CPA permeability and hydraulic conductivity within the BBB, due to the high concentration of aquaporin 4 in the cell membrane of astrocytic perivascular endfeet in orthogonal arrays of particles (55). We assume that during our slice immersion protocol, CPAs distribute via the freely accessible extracellular space (56), whereas during our *in situ* perfusion protocol, CPAs distribute through the brain tissue via the astrocyte, combined with a fluid efflux from the perivascular space. Most astrocytes interface at least one capillary and extend throughout the whole parenchyma with highly branched submicron processes (57). Shock-loading and interleaved equilibration should therefore induce significant volume excursions in astrocytes and the perivascular space relative to neurons and the endothelium, which do not express aquaporins, likely restricting water to passive transmembrane diffusion in the hypothermic state (55). This absence of aquaporins appears advantageous, as the neuronal cytoarchitecture withstands substitution of substantial proportions of its intracellular fluid with CPAs. A reduction of hydraulic conductivity in aquaporin-expressing cells like astrocytes may be achieved by knockdown (60, 61), sequestration (62, 63) or inhibition (64, 65). Circadian variation in the hydraulic conductivity of astro - cytic perivascular endfeet might need to be considered as well (58, 59), and targeting aquaporins appears as a promising avenue for cryopreservation in general.

Hypothermic-hyperosmotic tolerance limits for cultivated cells have been reported to extend to 1250 mOsm saline solutions, with a sharp viability drop-off beyond this range (61). Correspondingly, our results indicate that the hypothermic murine brain can recover from acute dehydration to below 70% total mass (**Fig. 4Aii**), but not from the dehydration to below 45% total mass observed during gradual CPA loading (**Fig. 4Ai**). This should caution against maximizing the extent of cerebral dehydration in cryopreservation protocols (62-64). The mechanical effects of dehydration might be exacerbated in brain tissue due to its intertwined and anisotropic cytoskeletal architectures (65), and by differential dehydration of astrocytes compared to other cell types in the brain. CPA washout after cerebral vitrification *in situ* remains challenging. Theoretically, CPA washout with highly viscous hyperoncotic solutions at low flow rates circumvents discrepancies between CPA and water permeability by transitioning from diffusion-limited to perfusion-limited substance exchange in the vasculature (60). The present whole-brain vitrification protocol via interleaved equilibration and hyperoncotic washout requires further optimization, due to the variability and magnitude of cerebral dehydration, and the relatively low success rate. More established techniques like aldehyde-stabilization might be an alternative (66). Although our electrophysiological results after *in situ* vitrification may reflect best-case scenarios due to a selection bias toward samples exhibiting electrophysiological activity, the observed near-physiological responses after rewarming indicate that the brain can tolerate vitrification procedures, encouraging further optimization.

A general limitation of our study is the short observation period of acute brain slices post-vitrification due to their natural deterioration after 10-15 hours. Further research is needed to extend this period, e.g., using organotypic slice cultures and to characterize the transcriptional and translational effects of vitrification.

In conclusion, we demonstrate that the brain is remarkably robust not only to near-complete shutdown by hypothermia, but even to a complete shutdown in the vitreous state at -196 °C. This reinforces the tenet of brain function being an emergent property of brain structure, and hints at the potential of life-suspending technologies (14-16).

## Acknowledgments

We thank Andrea Eichhorn, Elke Kretzschmar, Maria Schulte, Birgit Vogler and Holger Meixner for technical help.

## Funding

German Society of Cryobanks (AG, JW)

German Research Foundation grants SFB 1483, FOR 5534 (AG)

Interdisciplinary Center for Clinical Research Erlangen project J111 (AG), ELAN project P166-26514166 (EYA)

## Supplementary Materials

Supplementary Text

Figs. S1 to S11

Tables S1 to S4

Movies S1 to S12

<u>Suppl_Movies</u>

